# Analysis of the role of N-linked glycosylation in cell-surface expression, function and binding properties of SARS-CoV-2 receptor ACE2

**DOI:** 10.1101/2021.05.10.443532

**Authors:** Raymond Rowland, Alberto Brandariz-Nuñez

## Abstract

Human angiotensin I-converting enzyme 2 (hACE2) is a type-I transmembrane glycoprotein that serves as the major cell entry receptor for SARS-CoV and SARS-CoV-2. The viral spike (S) protein is required for attachment to ACE2 and subsequent virus-host cell membrane fusion. Previous work has demonstrated the presence of N-linked glycans in ACE2. N-glycosylation is implicated in many biological activities, including protein folding, protein activity, and cell surface expression of biomolecules. However, the contribution of N-glycosylation to ACE2 function is poorly understood. Here, we examined the role of N-glycosylation in the activity and localization of two species with different susceptibility to SARS-CoV-2 infection, porcine ACE2 (pACE2) and hACE2. The elimination of N-glycosylation by tunicamycin (TM) treatment or mutagenesis, showed that N-glycosylation is critical for the proper cell surface expression of ACE2 but not for its carboxiprotease activity. Furthermore, nonglycosylable ACE2 localized predominantly in the endoplasmic reticulum (ER) and not at the cell surface. Our data also revealed that binding of SARS-CoV and SARS-CoV-2 S protein to porcine or human ACE2 was not affected by deglycosylation of ACE2 or S proteins, suggesting that N-glycosylation plays no role in the interaction between SARS coronaviruses and the ACE2 receptor. Impairment of hACE2 N-glycosylation decreased cell to cell fusion mediated by SARS-CoV S protein but not SARS-CoV-2 S protein. Finally, we found that hACE2 N-glycosylation is required for an efficient viral entry of SARS-CoV/SARS-CoV-2 S pseudotyped viruses, which could be the result of low cell surface expression of the deglycosylated ACE2 receptor.

**Importance:** Elucidating the role of glycosylation in the virus-receptor interaction is important for the development of approaches that disrupt infection. In this study, we show that deglycosylation of both ACE2 and S had a minimal effect on the Spike-ACE2 interaction. In addition, we found that removal of N-glycans of ACE2 impaired its ability to support an efficient transduction of SARS-CoV and SARS-CoV-2 S pseudotyped viruses. Our data suggest that the role of deglycosylation of ACE2 on reducing infection is likely due to a reduced expression of the viral receptor on the cell surface. These findings offer insight into the glycan structure and function of ACE2, and potentially suggest that future antiviral therapies against coronaviruses and other coronavirus-related illnesses involving inhibition of ACE2 recruitment to the cell membrane could be developed.

## Introduction

Severe acute respiratory syndrome coronavirus-2 (SARS-CoV-2) is a highly transmissible betacoronavirus that emerged in 2019 and is responsible for the current pandemic (1–7). Outcomes of human infection range from asymptomatic infection to severe clinical disease (8, 9). Infection is frequently associated with severe acute respiratory syndrome (SARS) but may also trigger other responses leading to multiorgan failure and death (5-7, 10-12). Several vaccines against SARS-CoV-2, approved for, emergency use, are being administered to the global population. For now, vaccines are highly effective and are the most effective strategy for controlling the disease. Concerns about vaccine escape variants and the broad tropism of the virus requires the continued pursuit of a broad range of antiviral strategies (13–15).

SARS-Cov-2 infection begins with the binding of the virus spike (S) protein to the cell surface receptor, ACE2, which results in fusion of the viral and cell membranes, and viral entry (4, 16–19). Other cell surface molecules, such as heparin sulfate, may also participate in infection (20). ACE2 also serves as a receptor for SARS-CoV, identificated and isolated in 2002 (21). The S protein contains 22 N-glycosylation sites, which play important roles in immune evasion, protein conformation, and cell tropism (17-19, 22-24). One of the unique properties of the SARS-CoV-2 S protein is the presence of a furin-specific cleavage site located between the S1 and S2 subunits, which may assist in viral entry (25–27). In addittion, other host enzymes including TMPRSS2 (transmembrane protease, serine 2) might contribute to viral entry of the virus (16, 25–28).

Human ACE2 (hACE2) is a type-I transmembrane glycoprotein that catalyzes the hydrolysis of angiotensin II (a vasoconstrictor peptide) into angiotensin (1–7) (29). hACE2 is composed of extracellular, transmembrane, and cytosolic domains (30, 31). The receptor contains 7 N-linked glycosylation sites, at amino acid residues 53, 90, 103, 322, 432, 546 and 690 (Fig. 1A) (30, 32–36). The sugar residues were confirmed by glycosidase treatment and by glycomic and glycoproteomic analysis (32–36). The presence of a O-glycosylation site at T730 was also reported (34). One function that has been proposed for the N-linked glycans is a direct modulation of Spike-hACE2 binding (34–36). In particular, the glycans of hACE2 at N90, N322 and N546 are all reported to interact with the SARS-CoV-2 S protein (35, 36). Recent studies evaluated the contribution of hACE2 N-glycosylation in the interaction with SARS-CoV-2 viral S protein (37, 38). The structures of the sugars on the hACE2 receptor were modified genetically or enzymatically, generating different hACE2 glycoforms (37, 38). They concluded that S binding with ACE2 is slightly influenced by the N-linked glycans present in the hACE2 receptor and that the hACE2 glycans play no role in viral entry (37, 38). However, the N-linked sugars present on the Spike protein were critical for the virus to enter the host cells (38, 39).

**Figure 1.**
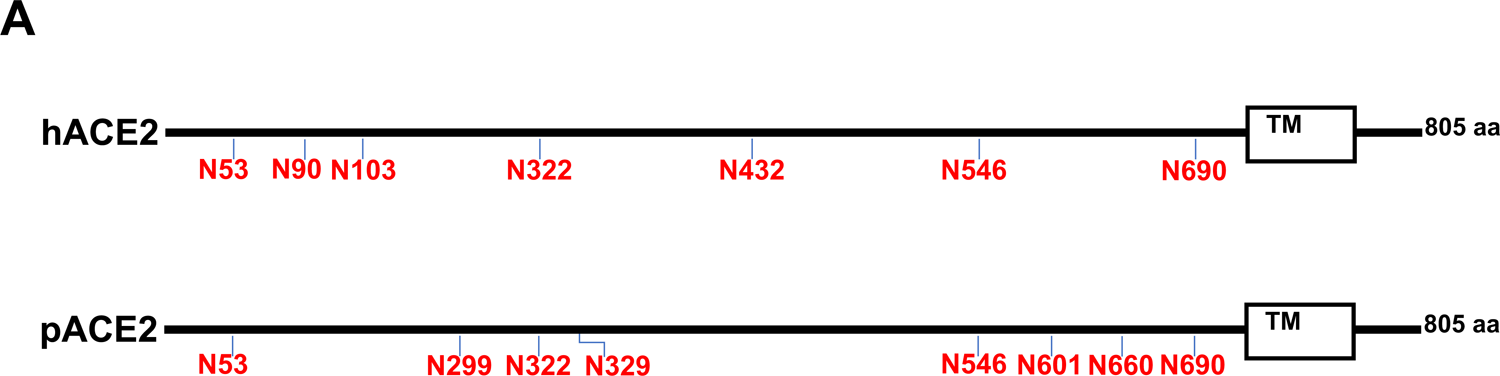
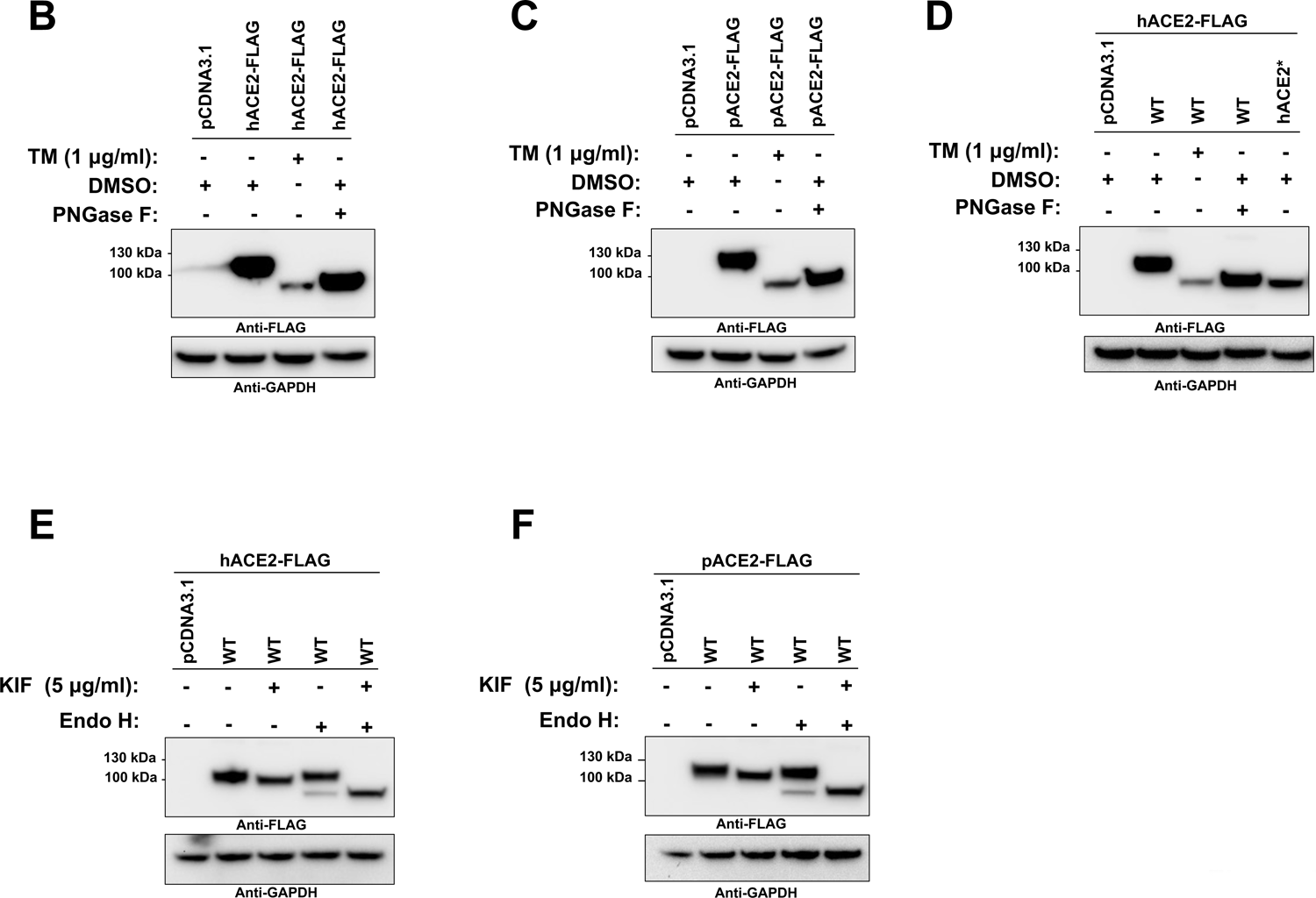
Effect of N-glycosylation inhibition on expression and electrophoretic mobility of hACE2 and pACE2. (A) Schematic representations of the N-glycosylation sites of ACE2 receptors from human and pig. The hACE2 N-linked glycosylation sites, --Asn-Xaa-Ser/Thr-(Xaa≠P)--, are indicated in red. 8 potential N-glycosylation sites in the pACE2 are also indicated in red. The box is the transmembrane domain. (B and C) HEK293T cells were transfected with plasmids expressing either hACE2 (B) or pACE2 (C) and treated with TM for 16 h before harvesting, or digested with PNGase F. ACE2 was detected with anti-FLAG antibody. (D) Western blot analysis of hACE2 and hACE2* expression levels in transfected HEK293T cells. Cells were treated with TM and cell lyastes digested with PNGase F. (E and F) Western blots of hACE2 and pACE2 expression levels after KIF treatment for 16 hours. Cell lyastes were digested with EndoH and immunoblotted with anti-FLAG antibody. In all experiments GAPDH was used as a loading control. hACE2* refers to N-glycosylation sites removed by replacing Asn with Gln.

ACE2 from different species posses different glycosylation patterns (34), which may influence the tropism of the virus. For instance, the mouse ACE2 receptor, which is not susceptible to SARS-CoV-2 infection (4, 40), only has three N-glycosylation sites that share similarities with hACE2 (34). On the other hand, the porcine ACE2 (pACE2) receptor has eight potential N-glycosylation sites and shares four similar sites with hACE2 (Fig. 1A). In vitro studies found that expression of pACE2 receptor on non-permissive cells to SARS-CoV-2 infection allows viral entry and infection (4, 40). In addition, there is growing evidence that pigs might be susceptible to SARS-CoV (41) or SARS-CoV-2 infection (42). Although, the results of these studies contradict previous reports indicating swine are not susceptible to SARS-CoV (43) or SARS-CoV-2 infections (44, 45). As part of this study, we compare the binding properties of hACE and pACE in response to deglycosylation.

It has been extensively proposed that N-glycosylation is involved in many important biological processes, including protein folding, enzimatic activity, trafficking and cell surface expression of proteins (46, 47). However, the importance of N-glycosylation for ACE2 function has not been previously investigated. This work explores the role of N-glycosylation of both pACE2 and hACE2 receptors in the cell surface expression, activity and in regulating direct Spike-ACE2 interactions. The results show that the total loss of glycans inhibits the proper cell surface expression of ACE2, but does not interfere with enzymatic activity. In addition, the complete removal of glycans from both S and ACE2 proteins does not inhibit their binding. Interestingly, hACE2 N-glycosylation decreased cell to cell fusion mediated by SARS-CoV S protein but not SARS-CoV-2 S protein-induced membrane fusion. Finally, we found that the presence of N-glycans in hACE2 is required for an efficient viral entry of SARS-CoV/SARS-CoV-2 S pseudotyped viruses, which can be attributed to the fact that deglycosylated ACE2 is less available in the cell surface.

## Results

### N-glycosylation inhibition induces accumulation of ACE2 in the ER

To understand the role of glycosylation in ACE2 function, we incorporated two N-glycosylation inhibitors tunicamycin (TM) and kifunensine (KIF). TM inhibits the first step of N-glycan biosynthesis, which results in the complete absence of glycan residues (48), while KIF is an inhibitor of ER-located mannosidase-I and complex N-glycosylation, resulting in the production of glycoproteins lacking the characteristic terminal sugar found on mature N-glycans (48, 49). Following transfection of 293T cells with a hACE2-expressing plasmid and treatment with TM, immunoblot analysis of whole cell lysates was performed. Treatment of the cells with TM resulted in a faster electrophoretic mobility shift for the hACE2 band showing the loss of N-linked glycosylation (Fig. 1B, lane 3). Digestion of untreated cell lysates with PNGase F, which cleaves all N-glycans, resulted in a hACE2 band similar in size to that of TM-treated cells, confirming the deglycosylation of hACE2 (Fig. 1B, lane 4). Similar results were obtained when we performed the same experiments with pACE2 (Fig. 1C). Next, to further confirm that TM treatment and PNGase F digestion produced a N-glycosylation-deficient hACE2, we generated a hACE2 variant, in which all the N present in the consensus N-glycosylation sites were replaced by Q (for clarity, this new mutant is herein referred to as hACE2*). As expected, expression of hACE2* in 293T cells resulted in detection of a band with a similar molecular weight to that of TM-treated lysates or samples digested with PNGase F (Fig. 1D; lane 5). Interestingly, TM treatment showed reduced levels of ACE2 (Fig. 1B, 1C and 1D). The effect of TM was mimicked by hACE2* construct, which lacked glycans. Together, these data suggests that core N-glycosylation is important for ACE2 biosynthesis.

KIF treatment of transfected cells did not have significant impact in the electrophoretic mobility of either hACE2 or pACE2 and generated bands with a slighy shorter molecular weight (Fig. 1E and 1F; lane 3). However, the digestion of KIF-treated cell lysates with Endo H, which cleaves only the high-mannose and hybrid branches of N-glycans, resulted in bands at a lower molecular weight compared to untreated controls (Fig. 1E and 1F; lane 5). In addition, untreated cell lysates digested with Endo H generated two different bands for both ACE2 proteins, one band with a similar molecular weight as the control and a faint band with the same size as the KIF-treated cells (Fig. 1E and 1F; lane 4). These last results confirmed a small presence of high mannose/hybrid type glycans on both ACE2 receptors. Taken together, these results suggest that ACE2 contains complex-form and high-mannose N-linked structures, which are consistent with previous studies (32–35).

Since blocking of N-glycosylation might cause accumulation of unglycosylated proteins in the ER (46, 47), we tested the effect of N-glycosylation inhibition on the subcellular localization of ACE2. For this experiment, we studied the localization of ACE2 by immunofluoresnce in the presence of TM or KIF. As shown in Fig. 2A, TM treatment induced colocalization of human and procine ACE2 proteins with a ER marker, PDI (50, 51). In the absence of TM, ACE2 proteins were localized mostly in the cell surface and showed no accumulation in the ER (Fig. 2A). Similarly, incubation of the cells with the mannosidase-I inhibitor, KIF, had little or no effect on the cell-surface localization of ACE2 (Fig. 2B). These observations indicate that the mannosidase I activity, which is required for processing newly formed high-mannose glycoproteins in the ER into mature glycoproteins containing highly hybrid complex-type glycans (46), is not neccesary for the cell-surface expression of ACE2. Quantification of the colocalization between the ACE2 proteins and the ER marker verified the accumulation of the ACE2 in the ER after TM treatment (Fig. 2C).

**Figure 2.**
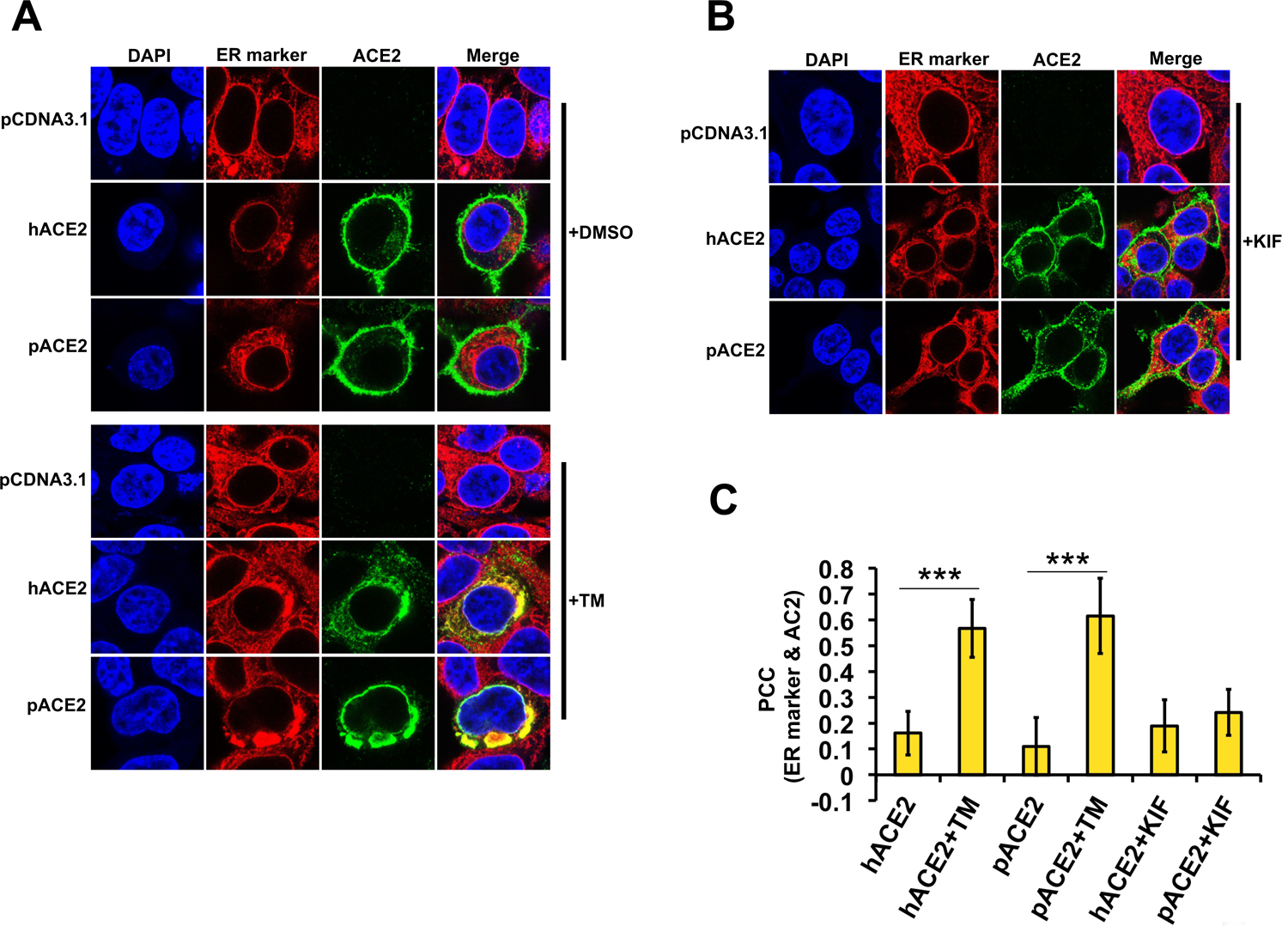
Inhibition of cellular N-glycosylation induces co-localization of ACE2 with the ER. (A) Semiconfluent monolayers of HEK292T cells grown on coverslips were transfected with either hACE2 or pACE2-expressing plasmids, and the cells were incubated with TM (1 μg/ml) (+TM) dissolved in DMSO or DMSO only for 16h, fixed and stained with anti-FLAG antibody and then with Alexa 488-goat anti-rabbit IgG (green). ER was visualized by staining cells with PDI monoclonal antibody, followed by Alexa 594-goat anti-mouse IgG (red). Nuclei were counterstained with DAPI (blue). (B) Same as A, except cells were treated with KIF (5 μg/ml) (+KIF) for 16h. (C) Quantification of colocalization of ACE2 proteins and ER. Individual cells (50–60) in each sample were selected randomly and analyzed. Pearson’s correlation coefficient (PCC) values are means ± SD from one representative experiment performed in triplicate. ***p < 0.0005 (as determined by two-tailed Student’s t-test) with respect to hACE2 and pACE2.

To further confirm that the N-glycosylation-deficient ACE2 proteins were arrested in the ER, a Golgi marker, Golgin-97, was used. As shown in Fig. S1, TM treatment did not show an overlap with the Golgi-marker, confirming that the N-glycosylation-defective ACE2 proteins were trapped in the ER unable to progress down the normal secretory pathway through the Golgi apparatus. Collectively, our data suggest that N-glycosylation is critical for cell surface expression of ACE2.

### N-glycosylation-deficient hACE2 variants accumate in the ER

Since TM is nonspecific, affecting the glycosylation of all proteins, we repeated the same localization experiments by constructing ACE2 proteins lacking different glycosylation sites. Mutants were constructed by replacing N with a Q. Following transfection of 293T cells with the hACE2 mutant constructs, immunoblot analysis on whole cell lysates was performed. The highest molecular weight band was found in the wild-type protein. As the number of N-glycosylation sites decreased, the migration size was reduced (Fig. 3A). Next, we investigated the cellular distribution of the N-glycosylation-deficient mutants expressed in 293T cells by immunofluorescence. The results revealed that the triple and quadruple mutants were expressed mostly on the cell surface similar to the wild-type hACE2 (Fig. 3B and 3C). Colocalization of the quadruple mutant and ER was also observed in some cells (Fig. 3B and 3C), whereas the other mutants, in which most or all of the N-glysosylation sites were removed, strongly overlapped with the ER (Fig. 3B and 3C). These results indicate that the N-glycosylation-deficient hACE2 variants failed to exit in the ER and in turn were not expressed at the cell surface. To further verify that N-glycosylation-defective ACE2 variants accumulated in the ER, we next tested whether those variants colocalized with the ER chaprone, calnexin (52). ACE2 nonglycosylated forms and calnexin colocalized (Fig. 3D), confirming that N-glycosylation-deficient ACE2 variants accumulated in the ER. Finally, to further confirm that hACE2* is retained in ER, we analyzed its colocalization with Golgin-97. As shown in Fig. S1, hACE2* did not overlap with the Golgi-marker, confirming that the N-glycosylation-deficient hACE2 variant was retained in ER and unable to progress down the normal secretory pathway through the Golgi apparatus. Altogether, our results indicate that at least a partial N-glycosylation is necessary for the proper cell surface expression of hACE2.

**Figure 3.**
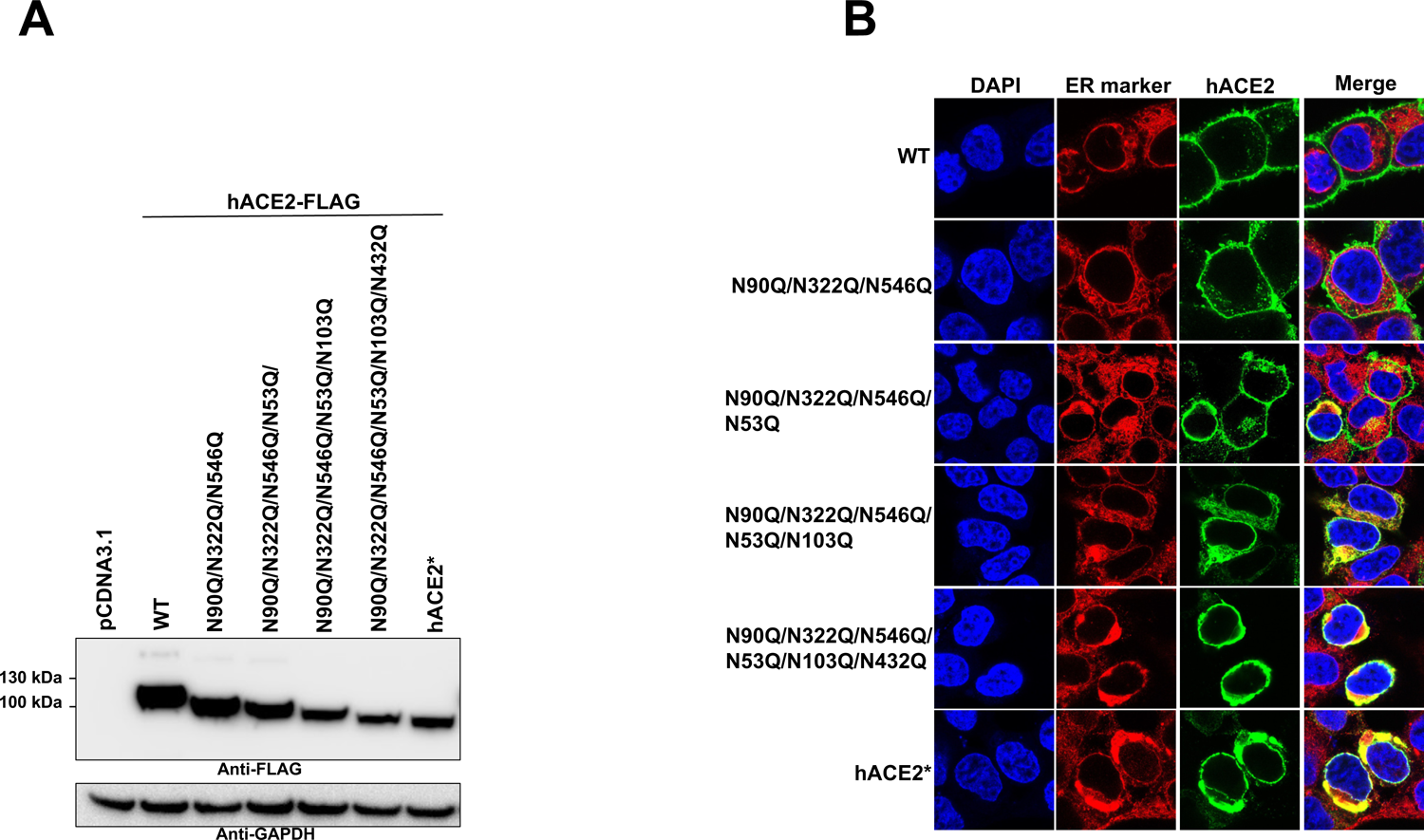
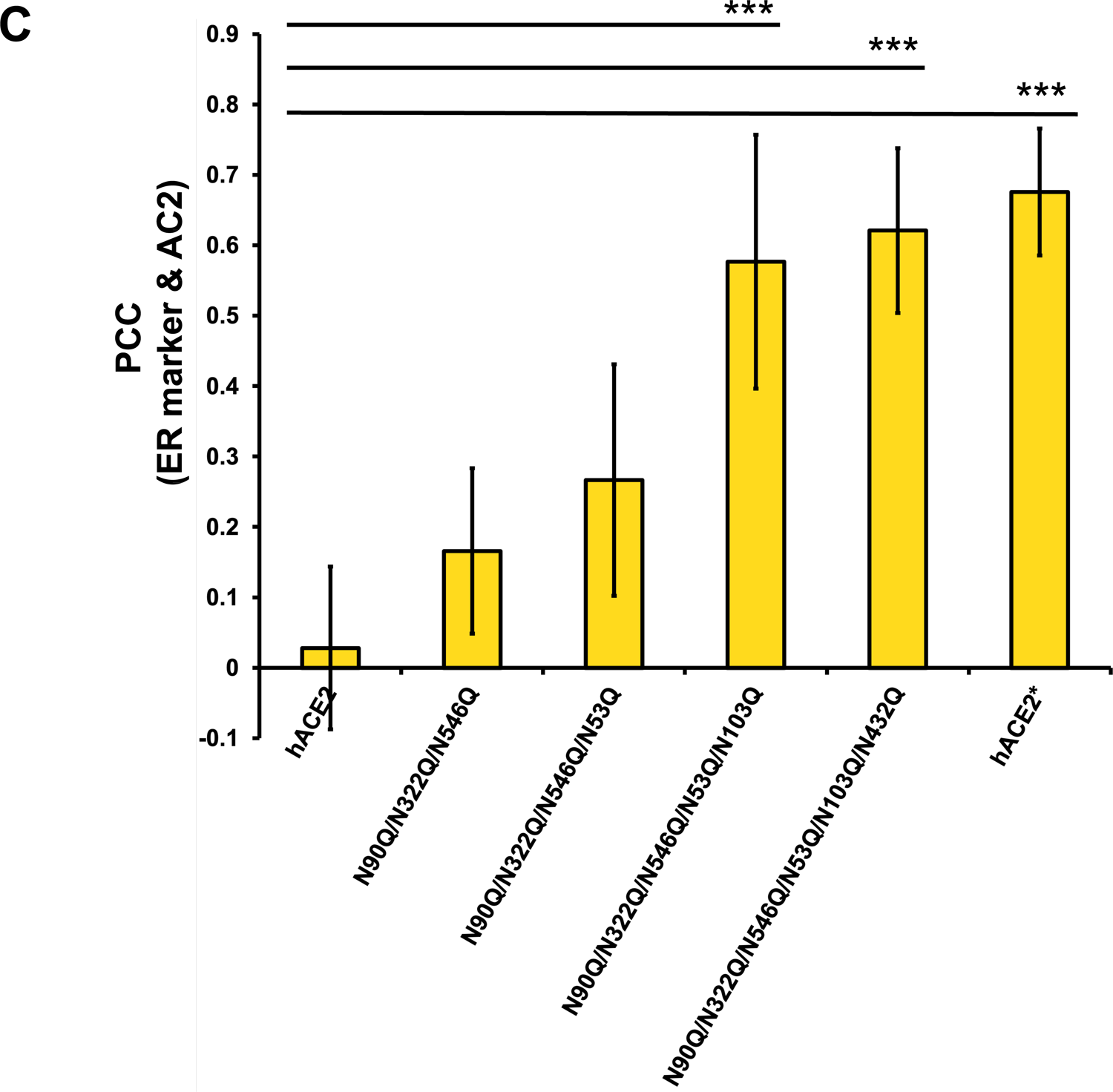
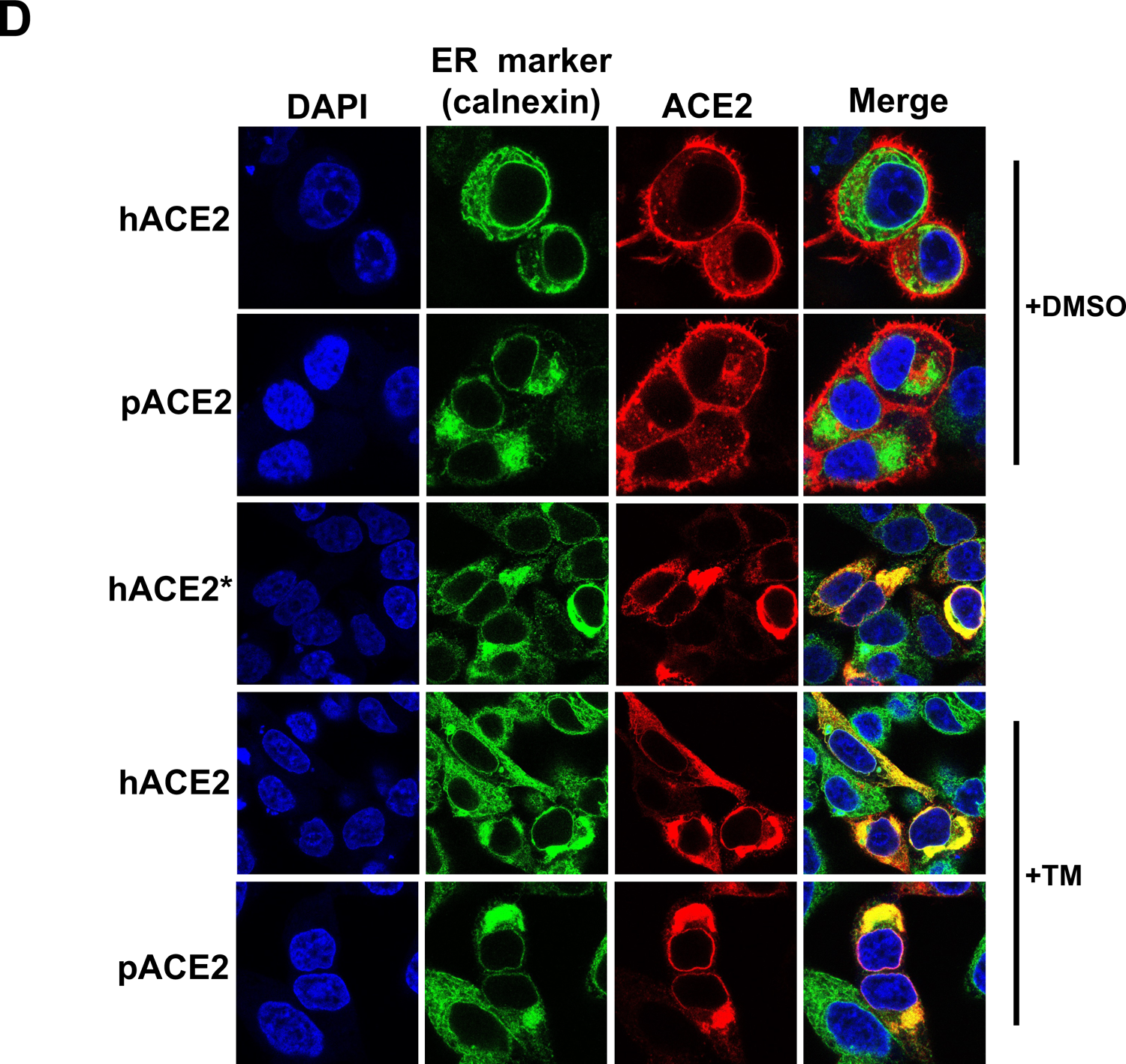
Colocalization of N-glycosylation-deficient hACE2 variants with ER. (A) Expression of the N-glycosylation hACE2 mutants was analyzed by Western blotting using anti-FLAG antibody and GAPDH as a loading control. Semiconfluent monolayers of HEK292T cells grown on coverslips were transfected with the indicated hACE2 mutants. All cells were fixed and immunostained as in Figure 2. (C) quantification of colocalization of ACE2 variants and ER was peformed as described in Figure 2. Pearson’s correlation coefficient (PCC) values between the ACE2 staining and intracellular endoplasmatic reticulum marker are shown on the plot. Values are means ± SD from one representative experiment performed in triplicate. ***p < 0.0005 (as determined by two-tailed Student’s t-test) compared with the wild-type hACE2. (D) Semiconfluent monolayers of HEK292T cells grown on coverslips were transfected with either hACE2, hACE2* or pACE2-expressing plasmids, and the cells were incubated with TM (1 μg/ml) (+TM) dissolved in DMSO or DMSO only for 16h, fixed and stained using anti-FLAG antibody and then with Alexa 594-conjugated goat anti-mouse IgG (red). ER was visualized by staining cells with calnexin antibody, followed by Alexa 488-conjugated goat anti-rabbit IgG (green). Nuclei were counterstained with DAPI (blue).

### N-glycosylation is critical for the proper cell surface expression of ACE2

The previous immunofluorescence experiments suggested that cell surface expression of the ACE2 nonglycosylated variants, generated either by TM treatment (Fig. 2) or by mutagenesis (Fig. 3), was reduced compared with wild-type ACE2. The abundance of nonglycosylated ACE2 or wild-type ACE2 on cell surface was analyzed by cell surface biotinylation. As shown in Fig. 4A and 4B, the surface density of both hACE2 and pACE2 proteins generated following TM treatment was significantly reduced compared with untreated cells. Similarly, the presence of hACE2* mutant on the cell surface was decreased compared with wild type hACE2 (Fig. 4C). Additionally, KIF treatment did not reduce the cell surface expression of both hACE2 and pACE2 (Fig. 4D), which is consistent with the immunofluorescence data shown above (Fig. 2C). In agreement with the findings shown in the previous section, these data demonstrate that ACE2 N-deglycosylation results in a pronounced decrease of cell surface expression.

**Figure 4.**
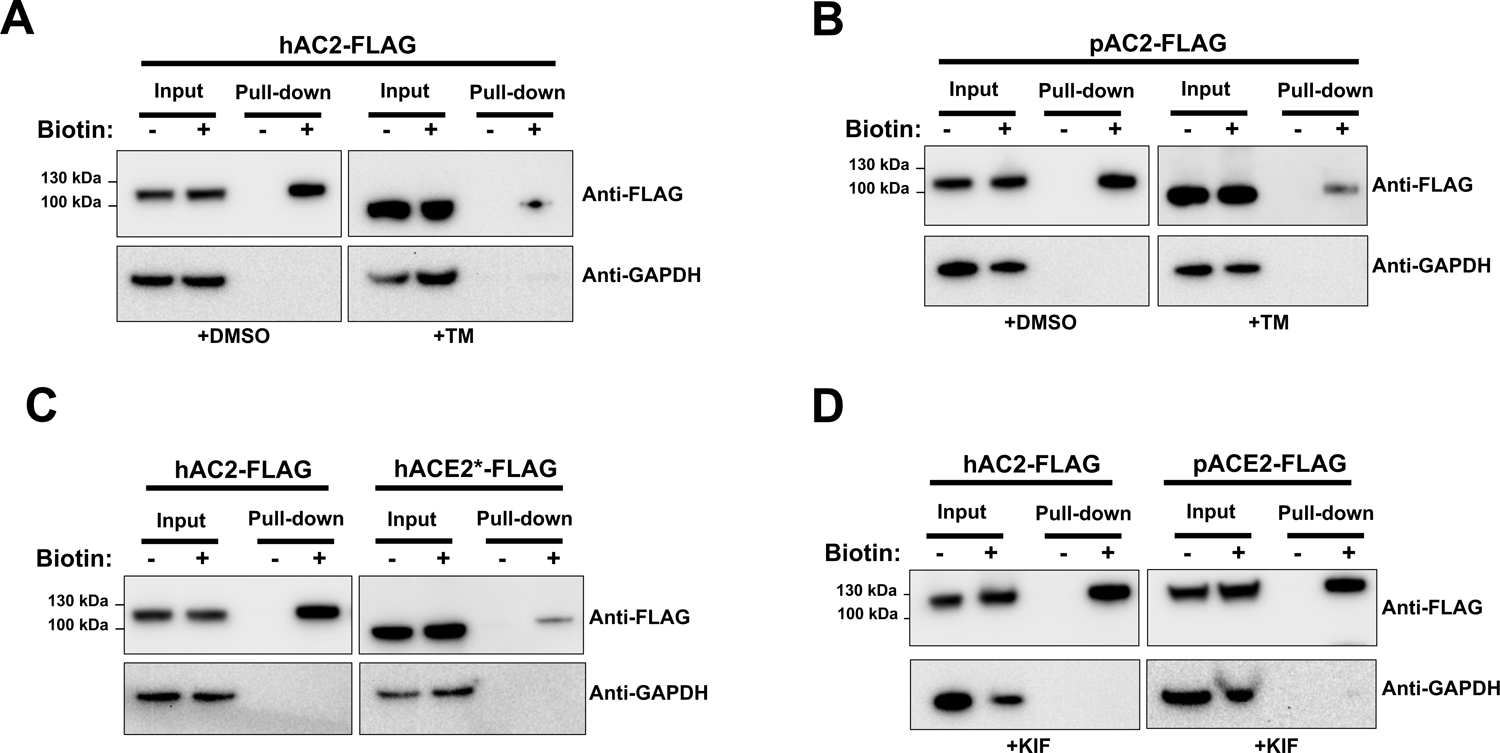
N-glycosylation inhibition and cell surface expression of ACE2. (A through D) HEK 293T cells were transiently transfected for 24 hrs with the indicated FLAG-tagged ACE2 constructs. Where stated, transfected cells were incubated with either TM (1 μg/ml) or KIF (5 μg/ml) for 16 h before biotin labelling. Cells were treated with (+) or without (-) biotin to label surface protein for 1 h at 4°C. After neutravidin agarose pull-down, proteins were immunoblotted with anti-FLAG antibody. GAPDH was used as the control to assess the purity of biotinylated plasma membrane fractions. Similar results were obtained in three independent experiments and a representative experiment is shown.

### N-glycosylation is not necessary for the carboxypeptidase activity of ACE2

To test the impact of N-glycosylation in the folding of ACE2 receptor, we determined the carbopeptidase activity of both nonglycosylated hACE2 and pACE2, generated by either TM treatment of cells or by mutagenesis. To directly analyze the carbopeptidase activity of ACE2 proteins, we tested the ability of immunoprecipitated ACE2 variants (Fig. 5A and 5B) to hydrolyze a synthetic peptide substrate. The ACE2 variants described in the western blot to the right of each graph were incubated with a fluorophore-labeled substrate. As shown in Fig. 5A and 5B, the constructs lacking glycosylation did not lose carboxippetidase activity when compared to wild type. These results show that N-glycosylation is not required for ACE2 protease activity and suggest that the N-glycosylation-deficient ACE2 variants are in a native conformational state.

**Figure 5.**
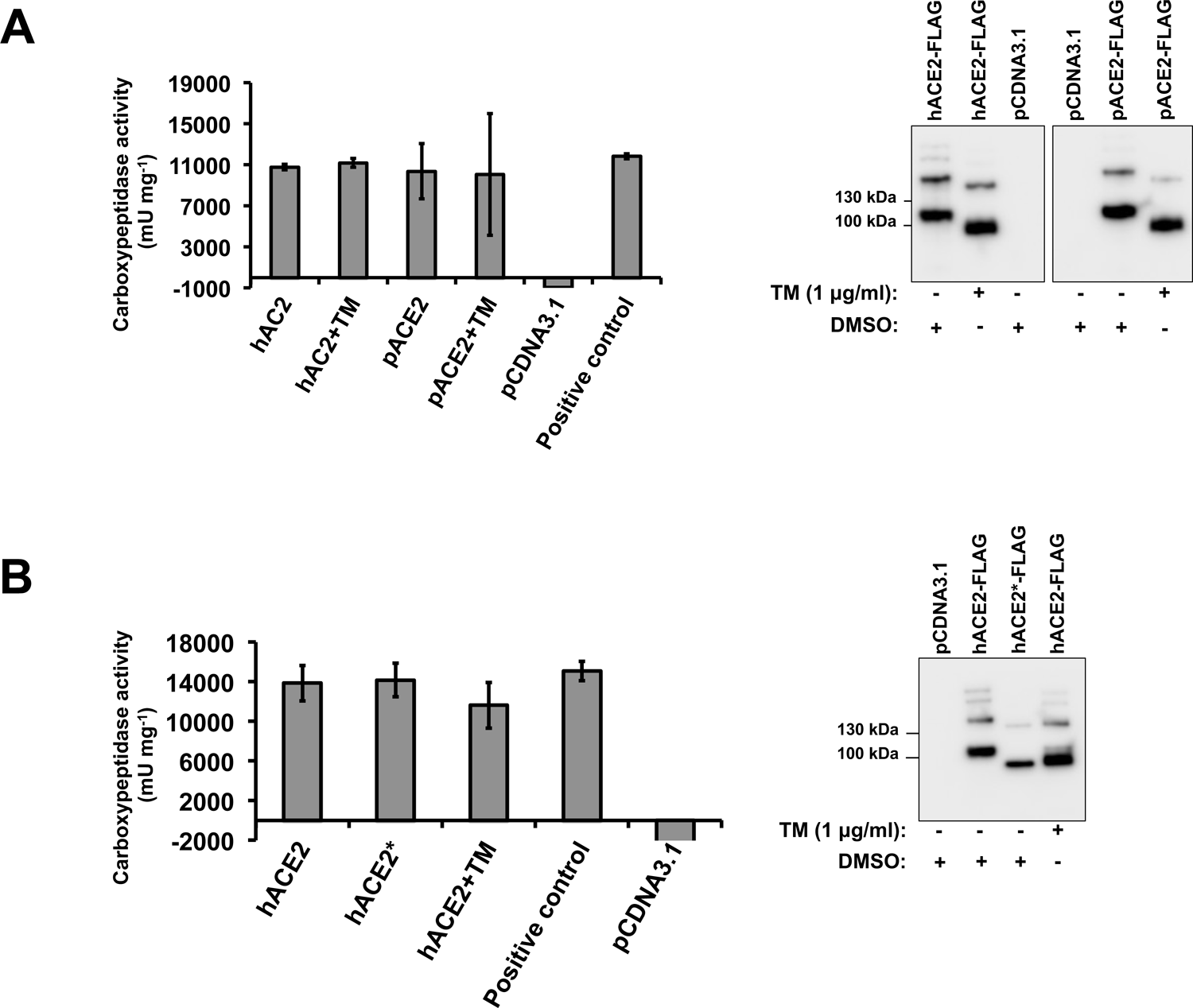
ACE2 Carboxypeptidase activity in the absence of glycosylation. (A) Plot showing the results of carboxypeptidase assays using the indicated immunoprecipitated FLAG-tagged ACE2 variants. A representative western blot with an anti-FLAG antibody that detects immunoprecipitated ACE2 variants is shown on the right. (B) the same as (A), but showing the results of carboxypeptidase assays using the immunoprecipitated FLAG-tagged hACE2* mutant. A representative western blot with an anti-FLAG antibody that detects immunoprecipitated hACE2 variants is shown on the right. Experiments were performed in triplicates and standard deviation is shown.

### N-glycosylation inhibition of ACE2 receptor and/or SARS-CoV-2/SARS-CoV-2 S protein does not disrupt S-ACE2 association

The ACE2 receptor is extensively glycosylated bearing high-mannose, hybrid, or complex carbohydrates distributed among its 7 N-glycosylation sites (Fig. 1A) (32–35). Several functions have been proposed for these N-glycans including a direct modulation of Spike-ACE2 binding (34–36). Recent studies investigated the role of ACE2 N-glycosylation in the interaction with SARS-CoV-2 S protein (37, 38). To this end, they generated different hACE2 glycoforms. Specifically, the N-linked sugars displayed by hACE2 were modified genetically or enzymatically (37, 38). However, the effect of a whole blocking of hACE2 N-glycosylation, by removing all the N-glycans from ACE2, on the binding to S protein was not investigated (37, 38). To assess the importance of N-glycosylation on ACE2-S binding, we analyzed the biochemical ability of N-glycosylation-deficient ACE2 variants containing a FLAG tag to interact with untagged S protein. For this purpose, we first independently transfected cells with plasmids expressing either hACE2 or S. Cells were lysed, and cell lysates containing FLAG-tagged hACE2 and untagged S protein were mixed. After precipitation with anti-FLAG beads, the eluted proteins by the FLAG peptide were separated by SDS-PAGE gels and analyzed by Western blotting using antibodies directed against the FLAG tag and the S protein. The SARS-CoV-2 S protein was efficiently coprecipitated by the anti-FLAG antibody, which is consistent with previous reports that demonstrated that hACE2 interacts with SARS-CoV-2 S protein (Fig. 6A) (4, 16–18, 26, 53–56). Similar results were obtained when we used pACE2 as the bait to pulldown the S protein (Fig. 6A). This result in agreement with the observation that expression of pACE2 on non-permissive cells to SARS-CoV-2 infection allows viral entry and infection (4, 40). Colocalization experiments confirmed the interaction of both hACE2 and pACE2 with SARS-CoV-2 S protein (Fig. 6B). Next, we tested the ability of N-glycosylation-deficient ACE2 variants, generated either by TM treatment or by mutagenesis, to bind S by performing similar methods. As shown in Fig. 7A and 7B, the nonglycosylated variants of both hACE2 and pACE2, generated after TM treatment, were able to interact with S protein. Similarly, the N-glycosylation-defective ACE2 mutant showed association with S protein (Fig. 7C). These results indicated that N-glycosylation is not required for the ability of ACE2 to bind SARS-CoV-2 S protein. Interestingly, a nonglycosylated variant of S generated after TM treatment, was able to interact with ACE2, suggesting that N-glycosylation of SARS-CoV-2 S protein is not required for its capacity to bind ACE2 receptor (Fig. 7A and 7B). It is important to point out that the expression of the nonglycosylated S variant was only detected in the pull-down assay when using ACE2 as bait (Fig. 7A and 7B), suggesting that the blocking of N-glycosylation might affect the stability or expression of the S protein. To confirm that the TM treatment generated a N-glycosylation-defective S variant, untreated S-containing lysates were digested with PNGase F. As expected, both treatments produced S protein bands of similar size (Fig. 7D). Consistently, N-deglycosylation of both ACE2 and S protein by mutagenesis or TM treatment, respectivally, did not disrupt their interaction (Fig. 7E). Additional colocalization data further confirmed the association between the N-glycosylation-deficient ACE2 variants and the Spike protein (Fig. 7F and 7G). Finally, we explored the ability of SARS-CoV S protein to bind nonglycosylated ACE2 variants. As shown in Fig. S2A and S2D, N-glysosylation-defective ACE2 variants from human and pig were capable to bind SARS-CoV S protein, which agrees with a previous work that showed that modifications of the N-linked glycan structure of hACE2 did not affect to its binding to the S protein (57). Colocalization results confirmed the S-ACE2 association (Fig. S2B and S2F). Moreover, a nonglycosylated S variant, generated by TM treatment, was able to interact with ACE2 receptors (Fig. S2A and S2D). Similar to SARS-CoV-2 S protein, the expression of the nonglycosylated S variant was only detected in the pull-down assay when using ACE2 as bait (Figure S2A and S2D), suggesting that the blocking of N-glycosylation might affect the stability or expression of the S protein. In addition, digestion of untreated S-containing lysates with PNGase F confirmed that TM treatment generated a N-glycosylation-defective S variant (Fig. S2C). Consistently, the N-deglycosylation of both ACE2 and S protein by mutagenesis or TM treatment of cells, respectivally, did not disrupt their interaction (Fig. S2E). Overall, these findings indicated that N-glycosylation is not required for the ability of ACE2 to bind either SARS-CoV S or SARS-CoV-2 S protein.

**Figure 6.**
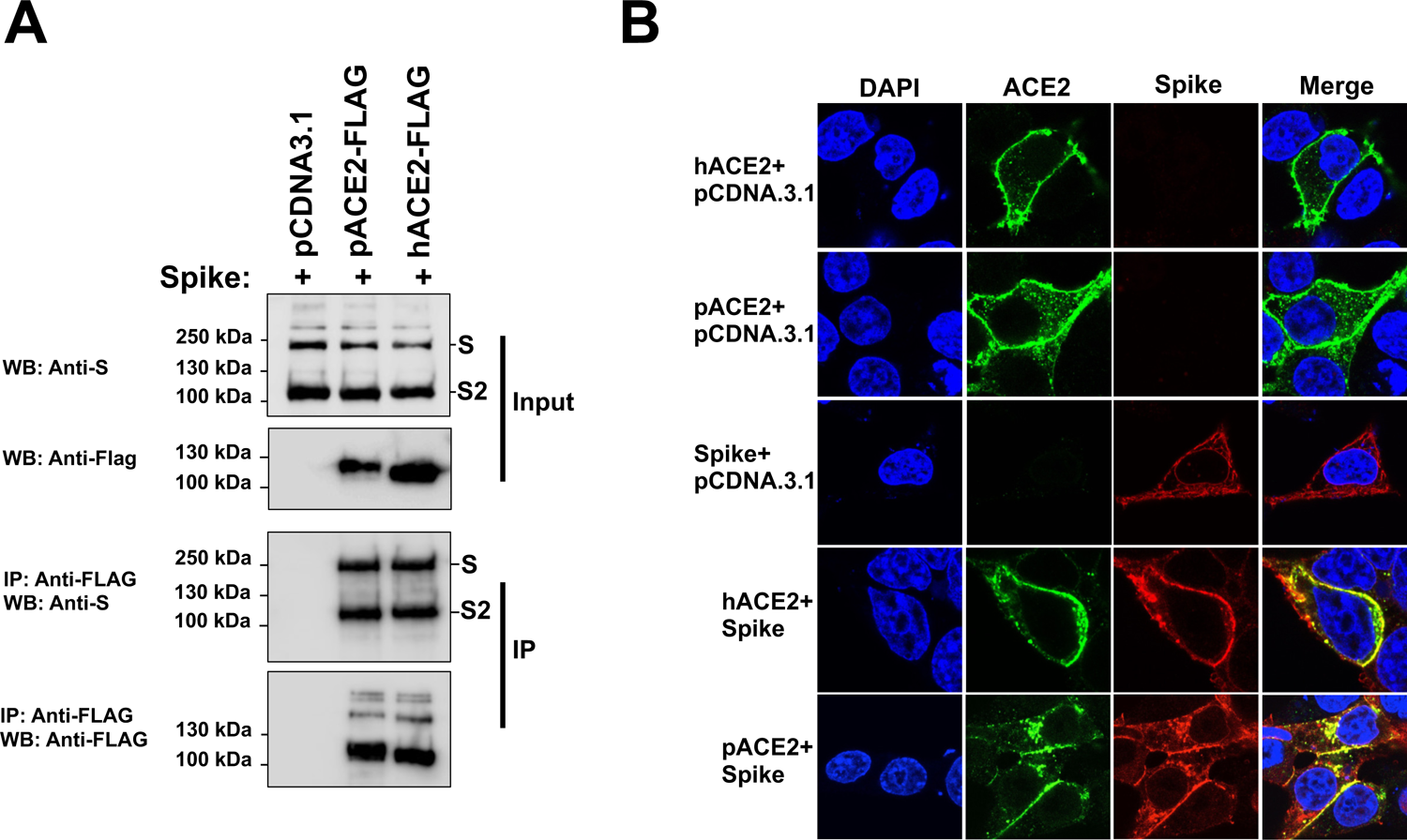
Co-immunoprecipitation assays and colocalization experiments to analyze ACE2-Spike interactions. (A) HEK 293T cells were independently transfected with vectors expressing the indicated FLAG-tagged hACE2 or pACE2 proteins. Other cells were transfected with a S-expressing vector. Cells expressing ACE2 variants or S protein were lysed at 24h post-transfection. The cell lysates were mixed in a 1:1 ratio, and incubated with anti-FLAG beads. To control for background binding of S protein to anti-FLAG beads, we performed similar experiments with HEK293T cells that were independently transfected with a S-expressing plasmid or an empty pCDNA3.1 vector. The amount of untagged and FLAG-tagged proteins in the lysates (Input) and immunoprecipitates (IP) was analyzed by Western blotting with anti-S and anti-FLAG antibodies. WB, Western blot; IP, Immunoprecipitation. Similar results were obtained in three independent experiments and representative data is shown. (B) Cellular co-localization of ACE2 variants and S protein. Cells were co-transfected with ACE2-expressing plasmids and a S-expressing vector. After 24 h, cells were fixed and immunostained using anti-FLAG antibody followed by Alexa 488-conjugated goat anti-rabbit IgG (green). The S protein was visualized using anti-S monoclonal antibody, followed by Alexa 594-goat anti-mouse IgG (red). Nuclei were counterstained with DAPI (blue). Similar results were obtained in three separate experiments and representative data is shown.

**Figure 7.**
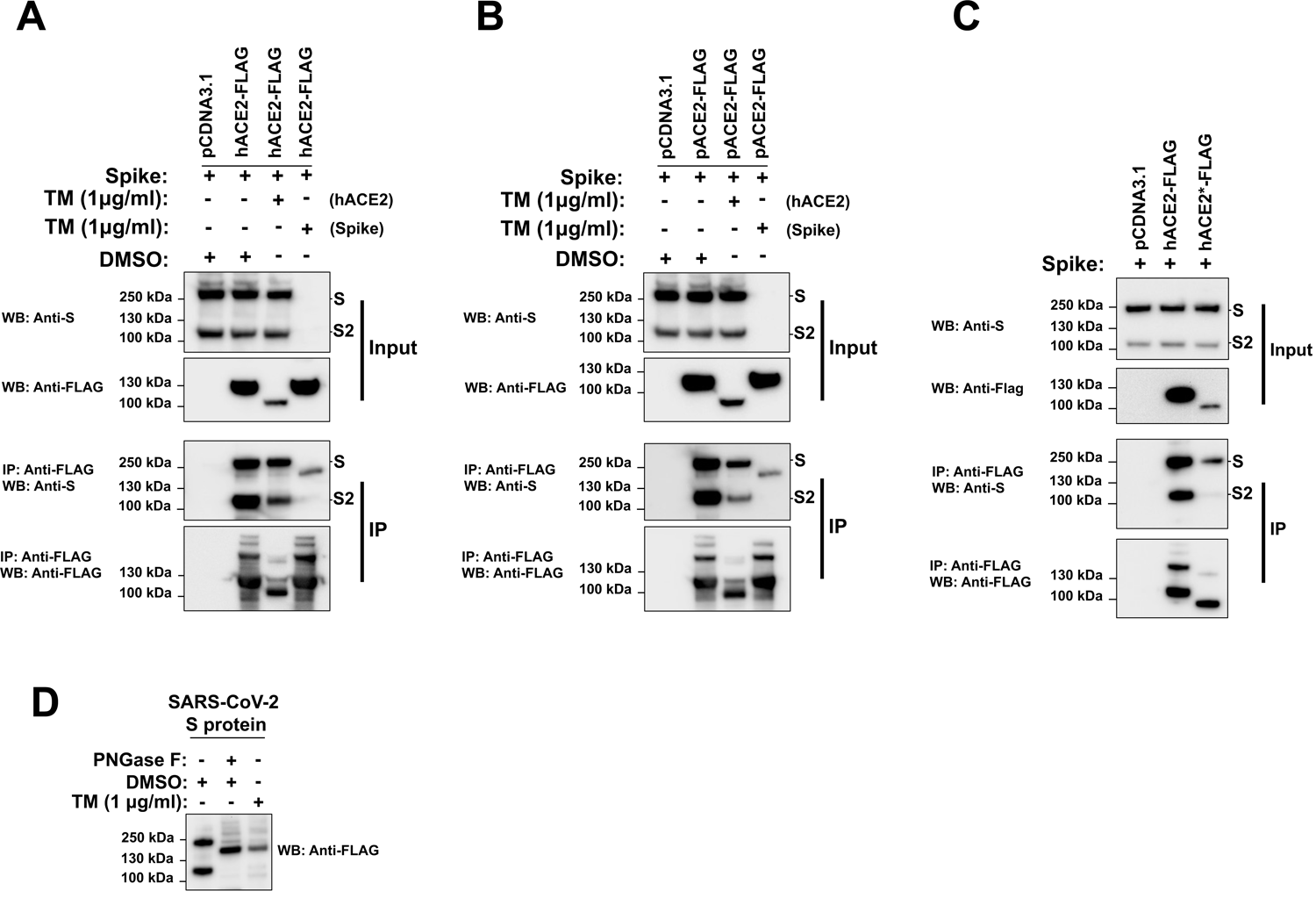
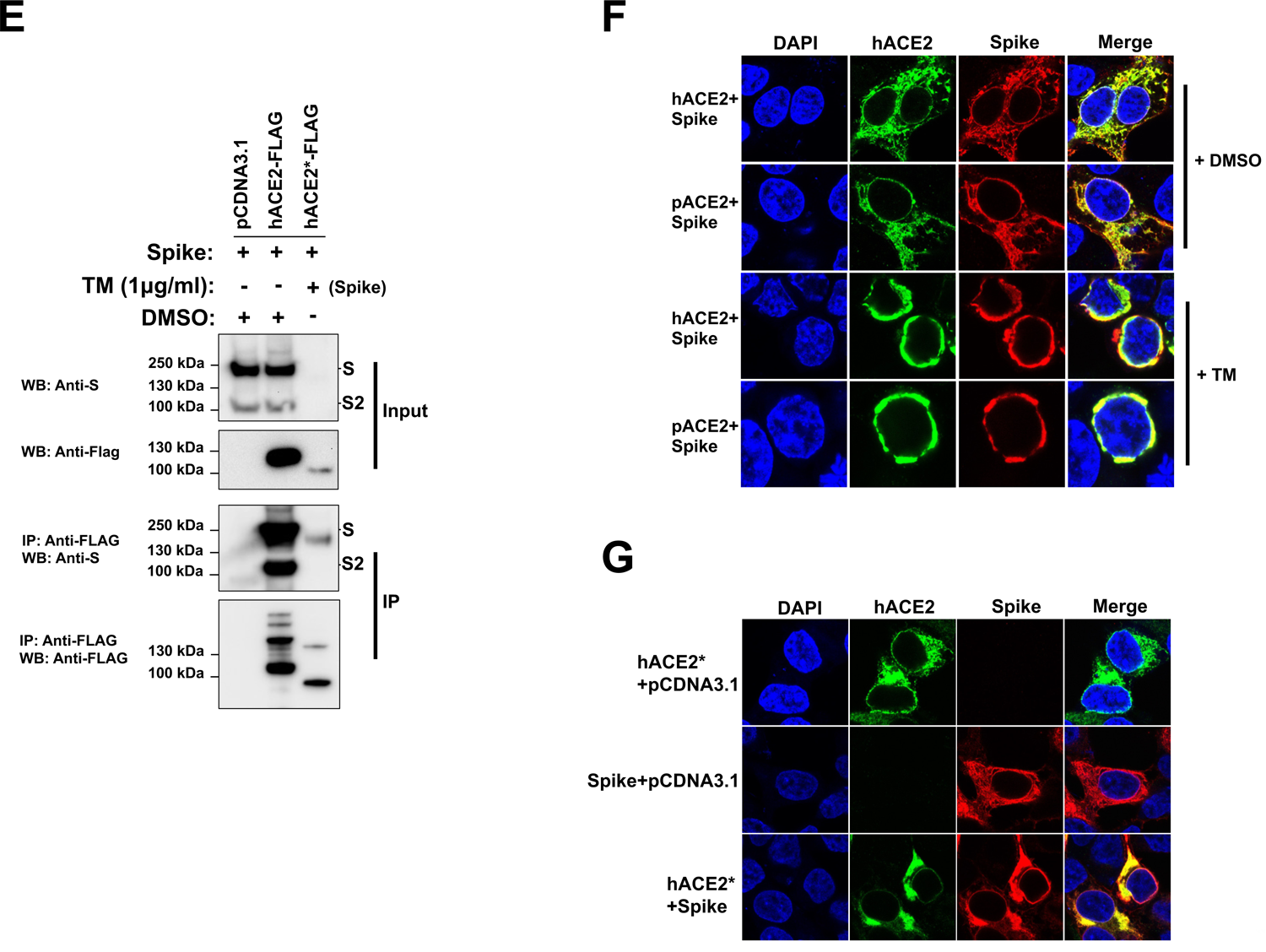
Role N-glycosylation in ACE2/S colocalization and binding (A through C) The same Western blot and immunoprecipitation experiments in Fig. 6A were repeated, but in the presence of TM (1 ug/ml). Cells were lysed 24 h after transfection. (D) 293T cells were transfected with a plasmid expressing SARS-CoV-2 S protein and treated with TM (1 μg/ml). 16 h post-treatment, the S protein was precipitated by using ACE2 as a bait (lane 3), as indicated in Figure 6. Cell lysates were also digested with PNGase F (lane 2). All the samples were analyzed by Western blotting using anti-S monoclonal antibody. (E) same (C), but both S and ACE2 protein were deglycosylated. (F and G) Cells were co-transfected with ACE2-expressing plasmids and a S-expressing vector. Cells were either untreated (+DMSO) or incubated with TM (1μg/ml) (+TM) dissolved in DMSO for 16h. All cells were fixed and immunostained using anti-FLAG antibody and then with Alexa 488-goat anti-rabbit IgG (green). S protein was visualized by immunostaining it with anti-S monoclonal antibody, followed by Alexa 594-goat anti-mouse IgG (red). Nuclei were counterstained with DAPI (blue). Similar results were obtained in three separate experiments and representative data is shown.

### hACE2 N-glycosylation impairment decreases cell to cell fusion mediated by SARS-CoV S protein but not SARS-CoV-2 S protein

Both furin or type II membrane serine proteases (TMPRSS)-mediated cleavage can trigger the fusogenic ability of both SARS-CoV-2 S and SARS-CoV S proteins, inducing receptor-dependent syncytial formation (25-27, 58-62). To investigate the effect of ACE2 N-glycosylation of either SARS-CoV-2 or SARS-CoV S glycoprotein-driven cell-to-cell fusion, we performed a widely adopted S-mediated cell–cell fusion assay (59, 63). The assay is based on the expression of GFP in one of the effector cells. Fusion is evident by the movement of GFP into the non-GFP expressing partner cell resulting in the formation of multinucleated large green cells. For this experiment, we used 293T cells that expressed either SARS-CoV-2 or SARS-CoV S protein and GFP (293T/GFP/Spike) as the effector cells, and 293T cells expressing either wild-type ACE2 or a N-glycosylation-defective ACE2 variant as the target cells ((293T/hACE2) (293T/hACE2*)). For the SARS-CoV S-mediated cell–cell fusion assay, effector cells were detached with trypsin and overlaid on target cells co-expressing ACE2 variants and TMPRSS2 to facilitate the fusion mediated by the S protein. In the case of SARS-CoV-2 S protein-triggered membrane fusion assay, cells were detached with 1 mM EDTA and overlaid on target cells. Syncytia formation was assessed by fluorescence microscopy. After effector cells and target cells were cocultured for 24h, the formation of big syncytia was observed in wild-type hACE2-expressing target cells when either of SARS-CoV or SARS-CoV2 S protein was used (Fig. 8A and 8B). No fusion was observed for GFP-expressing effector cells without S-expression or target cells without ACE2-expression (Fig. 8A and 8B), which confirms that S-receptor engagement is required for the S-mediated viral fusion. Next, we analyzed the fusogenic capacity of SARS-CoV-2/SARS-CoV S protein in target cells expressing the N-glycosylation-deficient hACE2 mutant. As shown in Fig. 8A and 8B, SARS-CoV S protein lost the ability to mediate cell–cell fusion when hACE2 N-glycosylation was impaired. This finding is consistent with the fact that the N-glycosylation-defective hACE2 variant is less expressed in the cell surface compared to the wild-type hACE2, which indirectly would reduce the fusogenic capacity of the S protein. On the contrary, SARS-CoV-2 S protein did not lose the ability to mediate the cell–cell fusion under the same conditions. No statistically significant differences were observed in syncytia formation comparing nonglycosylated with wild-type hACE2 (Fig. 8A and 8B). Thus, in contrast to SARS-CoV S protein, SARS-CoV-2 S protein was much more effective in mediating cell-cell fusion into target cells expressing a N-glycosylation-deficient hACE2 variant. Overall, these findings suggested that the SARS-CoV-2 S protein has higher capacity to mediate membrane fusion compared to the SARS-CoV S protein. This is in agreement with previous reports showing that the increased ability to mediate cell-cell fusion of the SARS-CoV-2 S protein is likely due to the presence of a furin cleavage site in its sequence (27, 59, 63).

**Figure 8.**
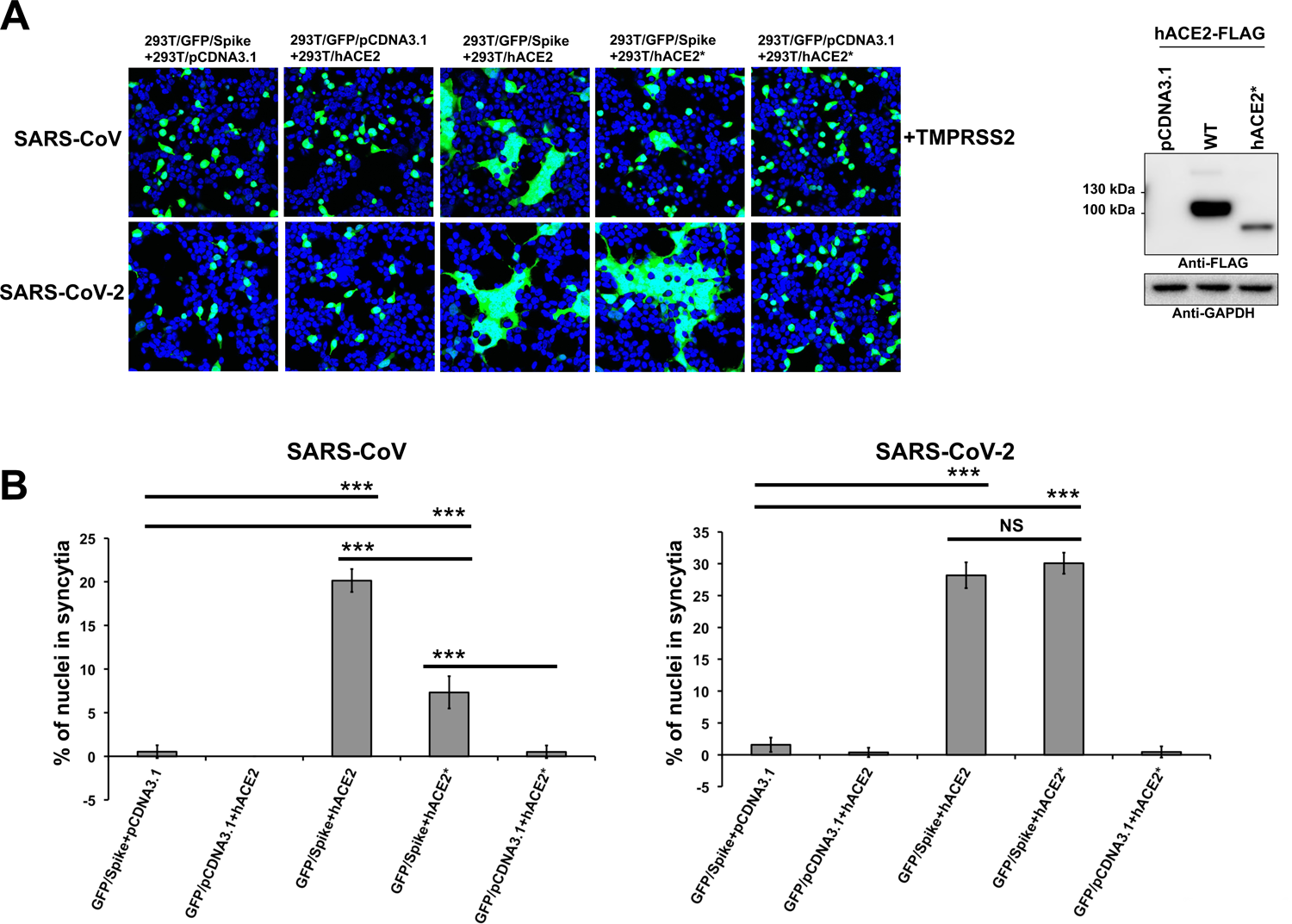
Effect of hACE2 N-glycosylation on SARS-CoV S and SARS-CoV-2 S protein fusion activity. (A) HEK 293T effector cells were co-transfected with a GFP-expressing plasmid along with one of the following plasmids: a plasmid expressing SARS-CoV-2 S protein, a plasmid expressing SARS-CoV S protein or an empty pCDNA3.1 plasmid. At 24 h post transfection, cells were detached and co-cultured with HEK293T target cells co-expressing either hACE2 or hACE2* and TMPRRSS2 for 24 h. Target cells transfected with an empty plasmid were included as negative control. Representative results are shown. A Western blot showing expression of hACE2 and hACE2* proteins is shown on the right. GAPDH was used as a loading control. (B) Quantitative representation of syncytia shown in (A). Values are means ± SD from one representative experiment performed in triplicate. ***p < 0.0005 (as determined by two-tailed Student’s t-test) with respect to the negative controls and hACE2*. NS, not significant.

### Effect of hACE2 N-glycosylation inhibition on the SARS-CoV/SARS-CoV-2 viral entry

Based on our previous findings demonstrating that N-glycosylation is required for a proper ACE2 cell surface expression, we hypothesized that viral entry of SARS-CoV-2 might be reduced in cells expressing a nonglycosilated variant of hACE2. To test this possibility, we investigated the impact of hACE2 N-glycosylation on viral entry. For this purpose, we determined whether GFP-expressing SARS-CoV-2 S pseudotyped viruses were able to transduce 293T cells expressing wild type hACE2 or a N-glycosylation-deficient ACE2 variant (hACE2*). The hACE2 proteins were transiently expressed (Fig. 9A) and then the ability of hACE2 receptors to allow SARS-CoV-2 S pseudovirions entry was tested. VSV-G pseudotyped viruses were used as a positive control. As expected, all transfected cells were effectively transduced by VSV-G pseudotyped viruses (Fig. 9B and 9C). Compared to cells transfected with empty plasmid (mock control), that was not susceptible to viral entry of SARS-CoV-2 S pseudotyped viruses, transduction of wild type hACE2-expressing cells with SARS-CoV-2 pseudoviral particles showed more GFP-positive cells. This observation is consistent with previous findings that demonstrated that hACE2 is the receptor of SARS-CoV-2 (4, 16, 17, 64). In contrast, the viral entry of SARS-CoV-2 S pseudotyped viruses in N-nonglycosylable ACE2-expressing cells was reduced compared to hACE2-expressing cells (Fig. 9B and 9C), suggesting that ACE2 needs to be N-glycosylated to support viral entry. Similarly, viral entry of SARS-CoV S pseudotyped viruses was also reduced in cells expressing a N-glycosylation-deficient variant (Fig. 9B and 9C). These results indicated that hACE2 N-glycosylation is required to allow an efficient viral entry of both SARS-CoV-2 and SASRS-CoV-2, which is in agreement with our previous results that showed that N-glycosylation is a prerequisite for the proper cell surface expression of hACE2.

**Figure 9.**
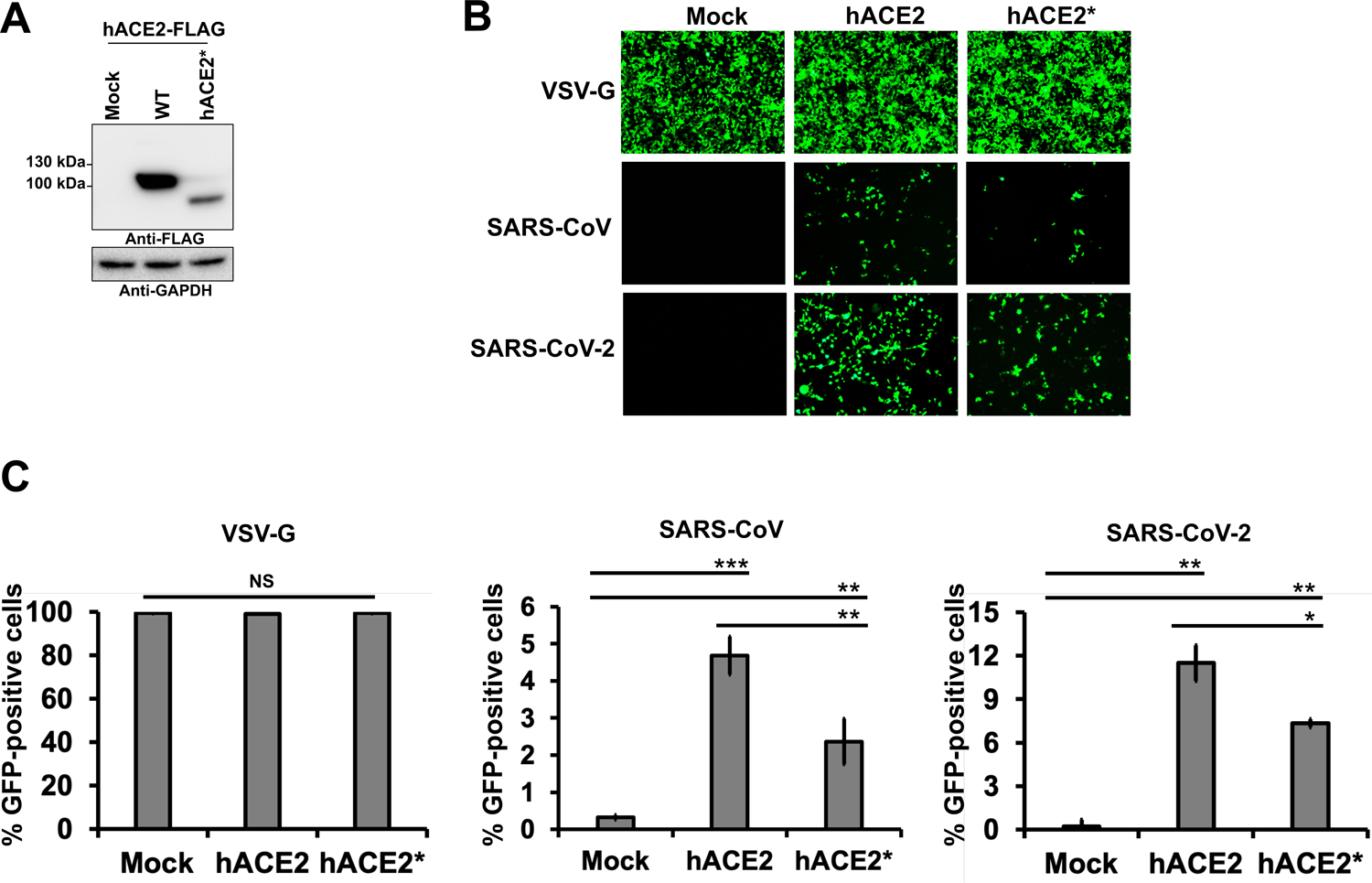
Contribution of ACE2 N-glycosylation to viral entry of SARS-CoV/SARS-CoV-2 S pseudotyped viruses. (A) HEK293T cells were transiently transfected with plasmids expressing hACE2 or hACE2* proteins, and their expression levels were analyzed by Western blotting using anti-FLAG and anti-GAPDH antibodies. (B) Representative fluorescence images of HEK293T cells expressing wild-type hACE2 or hACE2* after infection with equalised amounts of GFP-expressing SARS-CoV, SARS-CoV-2 or VSVG pseudotyped viruses. (C) The percentage of GFP-positive cells was measured at 72h post-infection by FACS. Wild-type VSV-G was used as a positive control. Values are means ± SD from one representative experiment performed in triplicate. *p < 0.05; **p < 0.005; ***p < 0.0005 (as determined by two-tailed Student’s t-test) with respect to wild-type hACE2 and Mock cells. NS, not significant.

## Discussion

Overall, the work presented here analyzes the importance of ACE2 N-linked glycosylation in cell membrane expression and protease activity as well as in various roles of ACE2, including binding to SARS-CoV-2/SARS-CoV-2 S protein, cell–cell fusion mediated by SARS-CoV/SARS-CoV-2 S protein and viral entry of SARS-CoV-2/SARS-CoV-2 S pseudotyped viruses. From these studies, we have learned the following: (1) N-glycosylation is a major determinant for the biosynthesis and the proper cell surface expression of ACE2, (2) ACE2 N-glycosylation is not required for its carboxipeptidase activity, (3) Association of SARS-CoV-2/SARS-CoV-2 S protein with ACE2 is not disrupted by N-glycosylation inhibition of ACE2 receptors or the viral proteins, (4), Impairment of hACE2 N-glycosylation affects cell–cell fusion mediated by SARS-CoV S protein but not the membrane fusion induced by SARS-CoV-2 S protein, and (5), hACE2 N-glycosylation is indirectly required for an efficient viral entry of the SARS-CoV/SARS-CoV-2 S pseudotyped viruses.

In agreement with previous studies (32–36), we verified, by digestion with glycosidases and by treatment with inhibitors that interfere at different stages of N-glycosylation biosynthesis, that hACE2 is N-linked glycosylated. Digestion with Endo H confirmed a small presence of high mannose/hybrid type glycans at hACE2 receptor. This finding is consistent with previous glycomic and glycoproteomic analysis that found that complex-type glycans were much more abundant than high mannose/hybrid type glycans across all N-glycosylation sites of hACE2 (34, 35). By using the same methods, we also found that pACE2, that showed four similar N-glycosyaltion sites with human (Fig. 1A), displays a similar N-glycosylation pattern as its homologous hACE2. These results suggest the presence of N-linked glycans in ACE2 receptors across different related species.

A large body of evidence describes important roles for N-glycosylation in protein stability and in cell surface expression of biomolecules (46, 47, 65-68). Our results demonstrate that the complete deglycosylation of ACE2 by site-directed mutagenesis or TM treatment results in a significant decrease in cell surface expression of the ACE2 receptor (Fig. 2 and 3). Reduced surface expression is the result of reduced protein expression (Fig. 1 and Fig. 4) along with increased retention in the ER (Fig. 2 and 3). As shown in Fig. 3B the loss of five N-glycosylation sites, represented by the N90Q/N322Q/N546Q/N53Q/N103Q is sufficient to retain ACE-2 in the ER. Colocalization studies showed that deglycosylated ACE2 was predominantly localized to the ER and not in the Golgi, suggesting that N-glycosylation is important for the biosynthetic processing from the ER. In the absence of N-glycans, Golgi processing is prevented, blocking the secretory pathway through the Golgi. Incubation of cells with KIF resulted in the loss of complex glycosylation of ACE2 but did not result in the retention in the ER. This result shows that complex N-glycosylation is not required for surface expression of ACE2 (Fig. 2 and 4), allowing the surface expression of ACE2 lacking complex sugars. This finding is consistent with a previous work that showed that inhibition of ER glucosidases with iminosugars did not affect ACE2 expression on the cell surface (57). These results show that the reduction of viral entry observed for the SARS-CoV/SARS-CoV-2 S pseudotyped viruses is likely the result of reduced expression of deglycosylated ACE2 on the cell surface.

Another potential role for N-glycosylation is the effect on the interaction between ACE-2 and the S protein. There are several examples of the requirement for N-glycosylation in formation of protein-protein interactions. For instance, N-glycosylation of the vasopressin V1a receptor is needed for optimal receptor-ligand binding (69) and N-glycosylation of P2Y12 receptor is necessary to trigger a proper downstream Gi-mediated signaling (70). In contrast, N-glycosylation is not required for the receptor functions of angiotensin II type-1 receptor (71), histamine H2 receptor (72) and the orphan G protein-coupled receptor Gpr176 (68). The interaction between the heavily glycosylated ACE2 and glycosylated S protein predicts a role for N-glycosylation in forming the receptor-S interaction. The published crystal structure and molecular modeling of hACE2, indicate that N90, N322 and N546-linked glycans interact with the SARS-Cov2 S protein (19, 35, 36, 73). Other studies show that the absence of N-linked glycans in hACE2 has a little or no effect on S binding (37, 38). Here we show that the complete removal of N-glycans by mutagenesis or TM treatment did not disrupt the interaction of both ACE2 receptors (pig and human) with SARS-CoV or SARS-CoV-2 S proteins. Moreover, removal of all N-glycans on ACE-2 and S proteins did not prevent co-precipitation, suggesting that glycans do not play a critical role in forming the interaction between ACE2 and S protein. In contrast, it has been reported for other viruses an important role of sialic acid present in the N-glycans on ligand-receptor interactions, including coronavirus like the Middle-East Respiratory Syndrome virus/MERS (74), parovirus (75) and influenza (76, 77).

In agreement with previous studies, we found that removal of N-glycans from hACE2 decreased the fusogenic ability of the viral S protein from SARS-CoV (Fig. 8A; (57)). One likely explanation for reduced fusogenic activity is the decreased surface expression of deglycosylated ACE2. Under the same experimental conditions, SARS-CoV-2 S protein did not lose the ability to mediate cell-to-cell membrane fusion. This observation is consitent with previous reports that showed that the higher ability to mediate cell-cell fusion of the SARS-CoV-2 S protein is likely due to the presence of a furin cleavage site in its sequence (27, 59, 63). In agreement with this notion, insertion of the S1/S2 furin-cleavage site significantly potenciated the capacity of SARS-CoV S protein to mediate cell–cell fusion but did not affect virion entry (78). In general, the S1/S2 furin-recognition site is missing in most of β-B coronaviruses, and their S proteins are uncleaved in normal conditions. For instance, in the case of SARS-CoV, that mainly uses the endosomal membrane fusion pathway to enter the host cell, its S protein is cleaved by endosomal cathepsin L and activated (79–81). The specific role of the S1/S2 cleavage site in the viral life-cycle of SARS-CoV-2 remains to be further investigated. A recent study showed that furin promotes both SARS-CoV-2 infectivity and cell-cell spread but it is not absolutely essential for SARS-CoV-2 infection and replication occurs in its absence, suggesting that furin-targeting drugs may reduce but not abolish viral spread (27). In line with these observations, a possible role of the furin cleavage site in reducing SARS-CoV-2 sensitivity to innate immune restriction was also proposed (82). Even though the role of the furin cleavage site in the SARS-CoV-2 life cycle remains unclear, in the this study we showed that it might function to overcome the low cell-surface expression of deglycosylated ACE2.

Our findings indicated that hACE2 N-glycosylation is required to allow an efficient viral entry of both SARS-CoV and SARS-CoV-2 virus, which might be atributted to the fact that N-glycosylation is necesary for the proper cell-surface expression of hACE2. The fact that fewer non-glycosylated ACE2 is available in the cell surface would imply less virus-receptor binding and therefore a decrease in the viral infection. Another posible explanation is that the removal of the N-glycans induced a misfolding of the hACE2 glycoprotein. However, our binding S-ACE2 experiments and protease activity studies suggested that N-glycosylation-deficient ACE2 variants are properly folded (Fig. 5, Fig.7 and Fig.S2). Alteration of N-linked glycans in ACE2 blocked its ability to support the transduction of SARS-CoV and human coronavirus NL63 (HCoV-NL63) S pseudotyped viruses by disruption of the viral S protein-induced membrane fusion (57). It remains to be determinated whether the reduced fusogenic activity is due to the aberrant glycan structure of ACE2 or to misfolding of the glycoprotein.

By analyzing the effect of blocking complex N-glycans formation of hACE2 receptor, it has been proposed that ACE2 glycans may not regulate viral entry of SARS-CoV-2 (38). In contrast to a full inhibition of N-glycosylation biosynthesis, our data showed that inhibition of complex N-glycosylation (by using KIF) did not alter the cell surface expression of ACE2. Moreover, a whole N-glycosylation depletion of the ACE2 receptor did not disrupt the S-ACE2 interaction, which would explain why the ACE2 N-glycans do not have a role in the viral entry of SARS-CoV-2. These obsevations support that idea that the reduction of the viral entry in cells expressing a non-glycosylated ACE2 is probably due to the lack of available cell-surface ACE2. However, the N-linked sugars present on the S protein were critical for the virus to enter the host cells since inhibition of complex N-glycan biosynthesis enhanced S protein proteolysis, suggesting that N-glycosylation might play a role regulating SARS-CoV-2 S protein stability (38, 39). In agreement with these findings, we observed that the expression of a nonglycosylated S variant was very low and only was detected by Western blotting when it was coprecipiated with ACE2, suggesting that the bloking of N-glycosylation might affect the stability or expression of the S protein.

Our data showed that the levels of wild-type hACE2 protein were higher than the N-glycosylation-deficient ACE2 mutant, suggesting that N-glycosylation inhbition might affect the stability or expression of the ACE2 receptor. Since N-glycosylation plays an important role in protein stability by protection against proteolysis (83–85), we investigated whether hACE2 protein stability is affected by the presence of its N-linked glycan motifs. Cycloheximide (CHX), an inhibitor of protein synthesis, was used to analyze the turnover rates of the hACE2 variants. We found that after up to 10h of drug treatment, hACE2 and hACE* levels did not change, suggesting tha N-glycosylation is not implicated in hACE2 stability and is not absolutely required for correct folding (Fig. S3). At this moment, our findings indicated that N-glycosylation is required for an efficient protein expression of ACE2. However, the exact role of N-linked glycosylation in the protein expression remains to be determined. Similarly, it was shown that the expression levels of other receptors such as rhodopsin (86), β2-adrenergic receptor (87), angiotensin II type-1 receptor (71) and the orphan G protein-coupled receptor Gpr176 (68) were all reduced by the depletion of N-glycosylation. However, this does not necessarily hold true for other receptors, for instance, it was shown that the expression levels of the orphan GPCR Gpr6 (88), α1-adrenergic receptor (89), M2 muscarinic receptor (90), histamine H2 receptor (72), vasopressin V2 receptor (91), PTH receptor (92), LH-RH receptor (93), and oxytocin receptor (94) were not significantly modified by blocking of N-glycosylation.

In conclusion, our data reveal an important role of the N-linked glycosylation in cell membrane expression of ACE2 but not in its protease actvity. Furthermore, our study demonstrates that N-deglycosylation of both ACE2 and S viral protein does not have a negative impact in their association. However, our findings also showed that depletion of ACE2 N-glycosylation recuded the viral entry of SARS-CoV and SARS-CoV-2, which is probably due to the lack of available cell-surface ACE2. These findings provide additional information regarding the glycan structure and function of ACE2, and potentially suggest that future antiviral therapies against coronaviruses involving inhibition of ACE2 traficking to the cell surface could be developed.

## Materials and Methods

### Plasmids

The plasmids, pCDNA3.1-hACE2-FLAG and pCDNA3.1-pACE2-FLAG, which express human ACE2 (GenBank accession number NM_021804.3) and pig ACE2 (GenBank accession number NM_001123070.1) fused to a FLAG epitope, respectively, were purchased from GenScript, as well as the pCDNA3.1-TMPRSS2-FLAG expressing TMPRSS2 (GenBank accession number NM_005656.4) fused to a FLAG epitope. pCDNA3.1-SARS2-Spike expressing full-lengh Spike (S) protein from SARS-CoV-2 (GenBank accession number QHD43416.1) was purchased from Addgene. Expression plasmid encoding S protein from SARS-CoV (GenBank accession number AAP13567.1) was purchased from SinoBiological. pCDNA3.1-SARS2-Spike Δ19 expressing a truncated SARS-CoV2 spike protein with the last 19 amino acids removed, was generated by using the Q5 Site-Directed Mutagenesis Kit (New England Biolabs), the plasmid pCDNA3.1-SARS2-Spike and the following mutagenic oligonucleotide primers: the sense primer was: 5’-CAGCTGCTGCTAGTTCGATGAGGACG-3’ and the antisense primer was 5’-CCGCAGGAGCAGCAGCCC-3’, following the manufacturer’s recommendations.

The pCDNA3.1-hACE2-FLAG plasmid and a Q5 Site-Directed Mutagenesis Kit (New England Biolabs) were used according to the manufacturer’s specifications to generate recombinant plasmids that expressed hACE2 N-glycosylation mutants, by substituting the N-linking site with Q. The following mutagenic oligonucleotide primers were used: for the production of mutation N53Q, the sense primer was 5’-TTATAACACCCAAATTACTGAAGAGAATG-3 and the antisense primer was 5’-TTCCAAGAAGCAAGTGAAC-3’; for the production of mutation N103Q, the sense primer was 5’-TCTTCAGCAACAAGGGTCTTCAGTGC-3’ and the antisense primer was 5-GCCTGCAGCTGAAGCTTG-3’; for the production of mutation N90Q, the sense primer was 5’-AGAAATTCAGCAGCTCACAGTCAAG-3’ and the antisense primer was 5’-TGTAGTGGATACATTTGG-3’; for the production of mutation N322Q, the sense primer was 5’-TGGTCTTCCTCAGATGACTCAAG-3**’** and the antisense primer was 5’-ACAGATACAAAGAACTTCTC-3’; for the production of mutation N546Q, the sense primer was 5’-TGACATCTCACAGTCTACAGAAGCTG-3’ and the antisense primer was 5’-CATTTGTGCAGAGGGCCT-3’; for the production of mutation N432Q, the sense primer was 5’-TCAAGAAGACCAAGAAACAGAAATAAACTTC-3’ and the antisense primer was 5’-AAATCGGGTGACAGAAGAC-3’; for the production of mutation N690Q the sense primer was 5’-TGCACCTAAACAAGTGTCTGATATC-3’ and the antisense primer was 5’-GTGACAAAGAAATTAAAGGAG-3’. The correctness of the constructs was confirmed by sequencing and by Western blot analysis of the expressed proteins. The hACE2* is a mutant in which the Asn (N) for all consensus N-glycosylation sites was replaced with a Gln (Q).

### Cells, antibodies and reagents

HEK 293T cells (ATCC CRL-3216 from American Type Culture Collection, Rockville, MD) and HEK 293T Lenti-X (Clontech/Takara Bio) were cultured at 37°C in 5% CO2 in Dulbecco’s modified Eagle’s medium (DMEM; Life Technologies) supplemented with 10% fetal bovine serum (FBS; Gibco), 2 mM L-glutamine (Life Technologies), and antibiotics (100 U/ml penicillin and 100 μg/ml streptomycin; Life Technologies).

The following reagents and their commercial sources were used: tunicamycin and cycloheximide (Sigma); Kifunensine (Tocris Bioscience); ACE2 activity kit (no. K897, BioVision); anti-FLAG-agarose beads (Sigma); and 3X FLAG Peptide (Sigma). The following antibodies were purchased from Thermo Fischer: SARS-CoV/SARS-CoV-2 Spike protein S2 monoclonal antibody (1A9), PDI mouse antibody (Clone: RL90), Alexa 594- and Alexa 488-conjugated antibodies against mouse and rabbit IgG, respectively. The monoclonal anti-FLAG M2 antibody was purchased from Sigma. The anti-GAPDH (D4C6R) monoclonal antibody, the Golgin-97 (D8P2K) rabbit monoclonal antibody, the Calnexin (C5C9) rabbit monoclonal antibody and the FLAG-tag (D6W5B) rabbit monoclonal antibody were from Cell Signaling.

### Transfection and immunofluorescence (IF) microscopy

Transfections of cell monolayers were done with the TransIT®-LT1 Transfection Reagent (Mirus) according to the manufacturer’s instructions (Mirus). Transfected cells were incubated at 37°C for 24-48 h unless otherwise stated. For indirect IF microscopy, cell monolayers grown on coverslips were transfected as indicated in the figure legends. At the indicated times, the monolayers were fixed with 4% paraformaldehyde. Fixed cells were permeabilized with 0.5% Triton X-100 in phosphate-buffered saline (PBS) and then blocked in PBS containing 2% bovine serum albumin. After incubation with primary antibodies for 1 h at RT or overnight at 4°C, the cells were incubated for 1 h with secondary antibodies and DAPI (4’,6’-diamidino-2-phenylindole). Images were obtained with a Nikon A1R laser scanning confocal microscope. The images were processed with NIS-elements software (Nikon).

### Quantitative analysis of fluorescent images

The fluorescent intensity of individual cells was measured by NIS-element software. The channel for each specific fluorescent signal was analyzed and the regions of interest (individual cells) were selected. The colocalization analysis was performed by using the colocalization analysis tool. In order to determine whether two fluorescent signals colocalized with each other, the Pearson’s correlation coefficient (PCC) values of individual cells were measured. Higher PCC values denoted higher levels of colocalization between the two fluorescent signals (See Fig. 2 and 3). Statistical analysis of the data was performed by using Student’s t test. p<0.05 was considered to be statistically significant.

### Western blot

Cellular proteins were extracted with a whole-cell extract buffer (50 mM Tris [pH 7.5], 150 mM NaCl, 0.5% Triton X-100, 10% glycerol, 1 mM EDTA, protease inhibitor cocktail [Sigma]). Cells lysates were resolved on 4–12% Bis-Tris NuPAGE gels (Invitrogen) and transferred to nitrocellulose membranes using a Trans-Blot Turbo Transfer System (Biorad). Protein bands were detected with specific antibodies using SuperSignal West Femto Maximum Sensitivity Substrate (Thermo Fisher) and viewed on a FluorChem R system (Proteinsimple).

### Immunoprecipitation assay

Human 293T cells were independently transfected with plasmids encoding FLAG-tagged mutant and wild-type hACE2 proteins, FLAG-tagged pACE2 protein and untagged SARS-CoV/SARS-CoV-2 S protein. For some experiments, cells were treated with tunicamycin (1μg/ml) for 16h before harvesting. After 24 h, the cells expressing each ACE2 variant or S protein were lysed in 1 ml of whole-cell extract buffer. Lysates were centrifuged at 14,000 rpm for 30 min at 4°C. Post-spin lysates were then precleared using protein A-agarose (Sigma) for 1 h at 4°C; a small aliquot of each of these lysates was stored as an input sample. Precleared lysates containing the differently tagged or untagged proteins were mixed in a 1/1 ratio and incubated with anti-FLAG-agarose beads (Sigma) overnight at 4°C to precipitate the FLAG-tagged proteins. Beads containing the immunoprecipitate were washed four times in whole-cell extract buffer. Subsequently, immune complexes were eluted using 200 μg of 3X FLAG Peptide /ml in whole-cell extract buffer without Triton X-100. The eluted samples were separated by SDS-PAGE and analyzed by Western blotting using anti-FLAG M2 monoclonal antibody or SARS-CoV/SARS-CoV-2 Spike protein S2 monoclonal antibody.

### Cell surface biotinylation assay

Human 293T cells were transiently transfected with plasmids encoding FLAG-tagged hACE2 variants or FLAG-tagged pACE2 protein for 24h. Where indiacted cells were treated with tunicamycin (1μg/ml) for 16h before harvesting. The cell surface biotinylation assay was performed using the Pierce Cell Surface Biotinylation and Isolation kit (Thermo Fisher) following the manufacturer’s recommendations. Briefly, the cell surface of 293T cells was biotinylated using EZ-Link sulfo-NHS-SS-biotin. Cells were lysed with whole-cell extract buffer and biotinylated proteins were recovered with Neutravidin beads. Input lysates and pulldown proteins were analyzed by western blotting with the anti-FLAG M2 monoclonal antibody (Sigma) and anti-GAPDH monoclonal antibody (Cell Signaling) as described above.

### ACE2 Activity Assay

Human 293T cells were transiently transfected with 10 µg of plasmid encoding FLAG-tagged hACE2 variants or FLAG-tagged pACE2 protein. Where indicated after transfection, cells were treated with tunicamycin (1μg/ml) for 16h before harvesting. At 24 h post-transfection, cells were lysed with whole-cell extract buffer and the FLAG tagged proteins were immunoprecipitated overnight at 4°C using anti-FLAG-agarose beads (Sigma), as described above. The immunoprecipitated extracts were processed in triplicates using the kit reagents and recommendations. The carboxypeptidase activity of the ACE2 variants was measured as fluorescence (Ex/Em = 320/420 nm) in kinetic mode using a Spectramax-ID5 plate reader (molecular devices) for 30 min to 2 h. A positive control was included by the kit. Immunoprecipitated extracts were also analyzed by western blot with the anti-FLAG M2 monoclonal antibody (Sigma) as described above.

### Flow Cytometry Analysis

Flow cytometry was performed to quantify the GFP-positive cells after SARS*-* CoV*-*2 or SARS*-*CoV S pseudotyped virus infection. Monolayers of 293T cells expressing hACE2 variants were infected for 48-72 h at 37°C with either SARS-CoV-2 or SARS-CoV S pseudotyped virus. Infection was performed in the presence of 8 µg/ml polybrene. HEK 293T cells transfected with an empty plasmid were used as the background negative control. After 48-72 hours post infection, cells were gently trypsinized (trypsin-EDTA). The percentage of GFP-positive cells was determined by flow cytometry (Cytek Aurora). 293T cells were initially analyzed by light scattering, where FSC (Forward Scatter) is a measure of size and SSC (side scatter) an indication of granularity and internal complexity. In all cases, a gate was applied on the bulk of the cells (95%), which excluded large aggregates. For each sample, 100,000-50,000 single cells were analyzed. To detect GFP, the samples were excited with a 488-nm laser coupled to an emission filter allowing the 515 nm wavelengths to go through. The data was processed with FlowJov10 software. All experiments were done in triplicate and repeated at least three times. Statistical analysis of the data was performed by using Student’s t test. p<0.05 was considered to be statistically significant.

### SARS-CoV-2 S pseudovirions production and viral entry

SARS-CoV-2 or SARS-CoV Spike-protein pseudovirions were generated by replacing the vesicular stomatitis virus (VSV-G) envelope protein of 3rd generation lentivirus with a mutant S protein possessing a deletion of 19 amino acid residues at the C-terminus. To generate these viruses, Lenti-X 293 T cells (Takara Bio) grown in 10 cm dish were transiently transfected with the following plasmids: 5 µg of pLenti-GFP (Cell Biolabs), 6 µg of psPAX2 and 0.9 µg of pCMV-VSVG (Cell Biolabs), or 0.9 µg of pCDNA3.1-SARS2-SpikeΔ19, or 0.9 µg of pCDNA3.1-SARS-Spike using the TransIT®-LT1 Transfection Reagent (Mirus) according to the manufacturer’s instructions. After overnight incubation, the medium was replaced with complete medium (DMEM + 10% FBS). At 48-72h after transfection, the supernatant was harvested, centrifuged at 800 × g for 5 min to remove cell debris and filtered through a 0.45 μm pore size PVDF syringe filter (Millipore), aliquoted and stored at −80°C until use.

In order to transduce cells with pseudovirions, virus was added either to mock transfected 293T or ACE2-transfected 293T cell lines at 24 hr post-transfection. Infection was performed in the presence of 8 µg/ml polybrene. After overnight incubation, cells were washed and returned to culture for 48 h and then subjected to flow cytometry analysis. GFP fluorescence was monitored daily. All flow cytometry and microscopy data presented in this study correspond to the 48-72 hr after transduction.

### Cell–cell fusion assay

HEK-293T cells expressing mutant or wild-type hACE2 variants were used as target cells. In some cases, the target cells were also transiently expressing type II membrane serine protease TMPRSS2. To prepare effector cells expressing S protein from either SARS-CoV or SARS-CoV-2, HEK-293T cells were co-transfected with plasmids encoding S glycoprotein and eGFP. For SARS*-*CoV S protein-mediated cell fusion, cells were lifted with trypsin (0.25%) at 24 h post transfection and overlaid on a monolayer of target cells at a ratio of approximately 1:1. For SARS*-*CoV-2 S protein-mediated cell fusion, cells were detached with 1 mM EDTA and overlaid on target cells at a similar ratio as above. After 24 h incubation, five fields were randomly selected in each well to count the number of nuclei in fused and unfused cells. Images of syncytia were obtained with a Nikon A1R laser scanning confocal microscope. All experiments were performed in triplicate and repeated at least three times. Statistical analysis of the data was performed by using Student’s t test. p<0.05 was considered to be statistically significant.

### Treatment with N-glycosylation processing inhibitors and glycosidases

Tunicamycin (TM) and Kifunensine (KIF) at a concentration of 1 μg/ml and 5 μg/ml, respectively, were used to inhibit N-glycosylation in transfected HEK-293T cells. Prior to treatment, cell lysates were mixed with glycoprotein denaturing buffer (New England Biolabs) and incubated at 100 °C for 10 min. To remove high-mannose oligosaccharides, cell lysates were treated with endoglycosidase H (Endo H) (New England BioLabs) for 1 h at 37°C. Peptide-N-glycosidase F (PNGase F) (New England BioLabs) was used to remove all N-linked oligosaccharides for 1 h at 37°C. Digested proteins were analyzed by Western blotting with anti-M2 FLAG monoclonal antibody (Sigma).

